# Structural alterations in the amygdala and impaired social incentive learning in a mouse model of a genetic variant associated with neurodevelopmental disorders

**DOI:** 10.1101/2023.06.14.545013

**Authors:** Takeshi Hiramoto, Akira Sumiyoshi, Risa Kato, Takahira Yamauchi, Gina Kang, Bailey Matsumura, Lucas J. Stevens, Rie Ryoke, Hiroi Nonaka, Akihiro Machida, Kensaku Nomoto, Kazutaka Mogi, Yukiko J. Hiroi, Takefumi Kikusui, Ryuta Kawashima, Noboru Hiroi

**Affiliations:** Department of Pharmacology, UT Health San Antonio, Texas 78229, USA; Institute of Development, Aging and Cancer, Tohoku University, 4-1 Seiryo-cho, Aoba-ku, Sendai 980-8575, Japan; National Institutes for Quantum and Radiological Science and Technology, 4-9-1 Anagawa, Inage-ku, Chiba 263-8555, Japan; Laboratory of Human-Animal Interaction and Reciprocity, Azabu University, 1-17-71 Fuchinobe, Chuo-ku, Sagamihara, Kanagawa 252-5201, Japan; Department of Cellular and Integrative Physiology, UT Health San Antonio, Texas 78229, USA; Department of Cell Systems and Anatomy, UT Health San Antonio, Texas 78229, USA

**Keywords:** *Tbx1*, volumetric MRI, social incentive motivation, 22q11.2 CNV, CNV, autism, schizophrenia, 22q11.2, amygdala

## Abstract

Copy number variants (CNVs) are robustly associated with psychiatric disorders and their dimensions and changes in brain structures and behavior. However, as CNVs contain many genes, the precise gene-phenotype relationship remains unclear. Although various volumetric alterations in the brains of 22q11.2 CNV carriers have been identified in humans and mouse models, it is unknown how the genes in the 22q11.2 region individually contribute to structural alterations and associated mental illnesses and their dimensions. Our previous studies have identified *Tbx1*, a T-box family transcription factor encoded in 22q11.2 CNV, as a driver gene for social interaction and communication, spatial and working memory, and cognitive flexibility. However, it remains unclear how *TBX1* impacts the volumes of various brain regions and their functionally linked behavioral dimensions. In this study, we used volumetric magnetic resonance imaging analysis to comprehensively evaluate brain region volumes in congenic *Tbx1* heterozygous mice. Our data show that the volumes of anterior and posterior portions of the amygdaloid complex and its surrounding cortical regions were reduced in *Tbx1* heterozygous mice. Moreover, we examined the behavioral consequences of an altered volume of the amygdala. *Tbx1* heterozygous mice were impaired for their ability to detect the incentive value of a social partner in a task that depends on the amygdala. Our findings identify the structural basis for a specific social dimension associated with loss-of-function variants of *TBX1* and 22q11.2 CNV.

## INTRODUCTION

A prerequisite to developing precision medicine in psychiatry is to identify risk factors, causative alterations in the brain, and their phenotypic targets in behavioral dimensions. Single nucleotide variants (SNVs) of single genes and copy number variants (CNVs) are robust risk factors for psychiatric disorders (1, 2, 3, 4, 5). How these risk factors manifest themselves as symptomatic elements of neurodevelopmental disorders is still poorly understood.

Evidence suggests that the shape and contour of the brain are robust constraints on behavioral functions (6). While whether this idea can be expanded into dysfunctions of the brain remains unclear, the ENIGMA (Enhancing NeuroImaging Genetics through Meta-Analysis) consortium has capitalized on a large sample size of idiopathic cases of mental illnesses and has identified many alterations in the volumes of the human brain regions. Individuals with schizophrenia have smaller volumes of the hippocampus, thalamus, amygdala, and nucleus accumbens; idiopathic autism spectrum disorder (ASD) cases are associated with smaller nucleus accumbens, amygdala, putamen, and pallidum (7, 8). Similar trends are seen in some CNV cases, but the precise volume changes vary among different CNVs (7, 8). The reproducibility of magnetic resonance imaging (MRI) data critically depends on the sample size (9, 10). It is difficult to collect a large sample of cases with rare variants such as CNVs and ultra-rare SNVs. Even when a large sample size is amassed by a consortium or in a database, a single outlier seems to inadvertently create an apparent overall significant difference between cases and controls (11).

Among CNVs, 22q11.2 CNV has been most intensively studied since the discovery of its association with mental illnesses in 1992 (12, 13). Data from consortia and meta-analyses of 22q11.2 CNV deletion cases show some consistent patterns: thicker cortical gray matter, but a reduction in focal thickness in the temporal and cingulate cortices and a reduced cortical surface area (14); decreased volumes of the hippocampus, thalamus, putamen, and amygdala, and increased volumes of the caudate nucleus and nucleus accumbens (7, 15). Among these volume changes, a volume reduction in subregions of the amygdala and a volume increase of the striatum are also seen in a mouse model of 22q11.2 deletion (16).

What remains unclear is how a large number of genes encoded in 22q11.2 CNV functionally contribute to volume alterations and consequent changes in behavior dimensions. Recent large-scale sequencing analyses have identified ultrarare cases of variants of CNV-encoded genes among non-CNV carriers with neurodevelopmental disorders (1, 2, 3, 4, 5). However, these ultrarare variants do not provide sufficient statistical power to establish them as causative genes, and sufficiently powered imaging studies are not feasible. Moreover, a mouse model of the whole 22q11.2 deletion is not suitable for identifying driver genes, as the impact of each gene is not isolated.

While mouse models of variants of small segments within CNVs and CNV-encoded single genes do not allow the determination of the CNVs’ functional roles in clinically defined disorders, they nonetheless provide a complementary means to identify potential genetic elements in dimensional aspects of psychiatric disorders (12, 17, 18). Human chromosome 22q11.2 CNV has been more comprehensively examined in mouse models compared with other CNVs, thanks to early studies that established its association with mental disorders (12, 13, 18, 19, 20). Such mouse studies have identified small chromosomal segments and single genes that impact specific dimensional aspects of intellectual disability, ASD, and schizophrenia (12, 17, 18).

Ultrarare variants of *TBX1* (which encodes T-box transcription factor 1) are found in individuals with ASD (21, 22, 23) and schizophrenia (5). Defective social interaction and cognition are common dimensions of idiopathic ASD and schizophrenia (19, 24, 25) and 22q11.2 hemizygosity carriers (26, 27). A dose alteration of *Tbx1* impairs social interaction and communication, spatial and working memory, and cognitive flexibility in mice (28, 29, 30, 31, 32). Moreover, *Tbx1* heterozygosity in mice alters the degree of myelination in the fimbria and cognitive speed (32), which is consistent with impaired speed of cognitive dimension in individuals with 22q11.2 hemizygosity (26, 33, 34).

We capitalized on this line of evidence to explore volumetric alterations and a behavioral dimension dependent on affected structures in congenic, constitutive *Tbx1* heterozygous (HT) mice. Our whole-brain volumetric MRI analysis determined that *Tbx1* heterozygosity is associated with smaller volumes of the amygdala and its surrounding regions and defective social incentive learning.

## MATERIALS AND METHODS

### Mice

Animal handling and use followed protocols approved by the Animal Care and Use Committee of the Albert Einstein College of Medicine, University of Texas Health Science Center at San Antonio, Tohoku University, and Azabu University in accordance with National Institutes of Health guidelines.

Our *Tbx1* HT mice were a congenic strain with a C57BL/6J background. The original non-congenic *Tbx1* HT mouse was backcrossed to C57BL/6J inbred mice for more than 10 generations to control the impact of uneven alleles originating from two strains on mutants and littermates (17). The genotype was determined using three primers: forward TTGGTGACGATCATCTCGGT and reverse ATGATCTCCGCCGTGTCTAG for detection of the WT genotype, and an additional reverse AGGTCCCTCGAAGAGGTTCA for detection of the HT genotype.

As no sex biases are seen in the prevalence of schizophrenia, psychosis, ASD (33, 35), or accuracy of social and cognitive dimensions (26) among carriers of 22q11.2 hemizygosity, we used either male or female mice for the various analyses.

### Brain sample preparation

We performed *ex vivo* MR scanning to achieve high resolution and a high signal-to-noise ratio due to the long scan time and use of a contrast agent (36). Using the standard procedure (37), 4-month-old female mice were anesthetized with pentobarbital (60 mg/kg, intraperitoneally) and transcardially perfused with 30 mL 0.01 M phosphate-buffered saline that contained 2 mM ProHance (Bracco Japan, Co., Ltd., Tokyo, Japan) and 1 mL/mL heparin (1000 USP units/mL). The mice were then perfused with 30 mL 4% paraformaldehyde (Wako, Tokyo, Japan) that contained 2 mM ProHance. The mice were decapitated, and the skin, lower jaw, ears, and cartilaginous nose tip were removed from the head. The skull structure, containing the brain tissue inside, was postfixed (4% paraformaldehyde and 2 mM ProHance) at 4°C overnight and then transferred to buffer (0.01 M phosphate-buffered saline, 0.02% sodium azide, and 2 mM ProHance) at 4°C overnight. The brains were placed in fresh buffer (0.01 M phosphate-buffered saline, 0.02% sodium azide, and 2 mM ProHance) at 4°C overnight. Immediately prior to scanning, ex vivo mouse brains were immersed in fomblin (MilliporeSigma, Burlington, MA, USA), a perfluorocarbon that minimizes susceptibility artifacts at the interface and limits sample dehydration during scanning.

### MR acquisition

MR scanning was carried out within 5 months after sampling (38). MRI data were acquired using a 7.0-T PharmaScan 70/16 system with a 23-mm-diameter birdcage Tx/Rx coil that was designed for the mouse brain (Bruker, Billerica, MA, USA) with the standard operational software (ParaVision 6.0.1; Bruker). Triplot images were acquired to ensure proper positioning of the sample with respect to the magnet isocenter. Shim gradients were adjusted using the MAPSHIM protocol with an ellipsoid reference volume covering the whole brain. T2-weighted images were obtained using a spin-echo 3D-RARE (rapid acquisition with relaxation enhancement) pulse sequence with the following parameters: repetition time = 250 ms, effective echo time = 32 ms, RARE factor = 4, field of view = 15 × 12 × 15.6 mm^3^, matrix size = 188 × 150 × 194, voxel size = 0.08 × 0.08 × 0.08 mm^3^, effective spectral bandwidth = 60 kHz, fat suppression = on, and number of averages = 16. The scanning time for T2-weighted images was approximately 8 hours for each mouse brain. As T1 tissue contrast between gray and white matter is less pronounced at a high magnetic field strength in rodents than in humans (39), we used T2-weighted contrast using our previously published procedure (40, 41). A voxel analysis was limited to the amygdala (small volume correction, FSL’s nonparametric permutation test, p < 0.05), independent of the atlas-based region.

### T2-weighted image analysis

Images were processed with the Advanced Normalization Tools software package (http://stnava.github.io/ANTs/) (42) in accordance with a published method (43): (i) images were reconstructed using ParaVision software, exported in DICOM format, and converted to NIfTI format using dcm2niix software; (ii) images were manually rotated and translated such that the origin of the coordinates occupied the midpoint of the anterior commissure to roughly match the standard reference space; (iii) a single reference image was manually skull-stripped using ITK-SNAP software (http://www.itksnap.org) to create the brain mask; (iv) other subject images were registered to the reference image and skull-stripped; (v) image nonuniformity was corrected, and image intensity was normalized; (vi) a minimum deformation template was constructed with the following SyN parameters: gradient step size = 0.1 voxels, update field variance = 3 voxels, and total field variance = 0.5 voxels (44); (vii) the obtained minimum deformation template was manually skull-stripped and registered to the atlas space (45); (viii) each region of interest volume was computed, divided by the total brain volume to account for the brain size differences, and statistically analyzed; (ix) the jacobian determinant for each mouse brain was computed, normalized by the total brain volume, and smoothed with a 0.4-mm full-width at half-maximum gaussian kernel; (x) voxel-based statistics were performed with permutation-based nonparametric testing with 500 random permutations as implemented in the FMRIB Software Library software package (https://fsl.fmrib.ox.ac.uk/fsl/fslwiki) (46). Brain regions were identified using the mouse brain atlas for MRI images (45, 47). In addition, each region of interest volume was converted to a mesh (VTK file format) using ITK-SNAP, and surface area was computed using ParaView software (https://www.paraview.org/).

### Immunohistochemistry

We used our published procedure (29, 48) to perform immunohistochemistry. Two- to 3.5-month-old female mice were anesthetized and perfused with physiological saline and then with 4% paraformaldehyde. The brains were kept in the fixative overnight at 4°C. Then, the mouse brains were cryoprotected in 20% glycerol in 0.1 M phosphate buffer overnight at 4°C. The mouse brains were cut into 40-μm thick sections. Free-floating sections were stained with a rabbit monoclonal anti-calretinin antibody (1:100, MA5-14540, Invitrogen/Thermo Fisher Scientific, Waltham, MA, USA) followed by a biotinylated goat anti-rabbit IgG (1:500, BA-2001, Vector Laboratories, Inc., Newark, CA, USA) and an avidin-biotin complex (Vectastain Elite, ABC kit peroxidase, Standard, Vector Laboratories, Inc, Newark, CA), using our standard 3,3′-diaminobenzidine staining procedure. Sections were examined using a Keyence BZ-X800 microscope (Keyence, Osaka, Japan). Each section containing the amygdalopiriform transition area was randomly chosen from both hemispheres. The calretinin-positive neuropil was delineated within the anteroposterior extent of the region (Bregma from −2.46 mm to −3.64 mm from bregma). The extent of stained neuropil was measured using Keyence analytical software. We used 12 images from 5 WT mice and 21 images from 4 HT mice. The average of each mouse was used for analysis.

### Social conditioned preference

Mice were maintained with their own parents until they were weaned at postnatal day 28. Mice were provided with food and water ad libitum. Each male mouse was then housed in a home cage with one male littermate: WT mice paired with a WT littermate; WT mice paired with a *Tbx1* HT littermate; *Tbx1* HT mice paired with a *Tbx1* HT littermate; *Tbx1* HT mice paired with a WT littermate. At 5–6 weeks of age, the experiment began, following a published procedure (49) with a slight modification. The test apparatus, made of opaque white polyvinyl chloride, had two compartments (each 170 × 170 × 170 mm) with paper bedding (Parumasu μ’, Material Research Center Co., Ltd., Japan) and Shepherd’s Cob bedding (Shepherd Specialty Papers, Richland, MI, USA). The two compartments were connected through a square opening (30 × 30 mm). The procedure included a preconditioning test session, two conditioning sessions with and without a social pair, and a postconditioning test session. During the preconditioning test session on Day 1, each mouse was allowed to freely move in the two compartments through the opening for 30 min. On Day 2, the opening was closed with a partition plate, and the mouse was confined with its home-cage littermate social partner to the compartment with paper bedding for 24 hrs. The mouse was then confined to the other compartment with Shepherd’s Cob bedding for 24 hrs (Day 3) without a partner. Immediately after the second conditioning, the test mouse was allowed to freely move between the two compartments through the open door (postconditioning session, Day 4). As a control for spontaneous preference change due to repeated testing, another set of mice was tested in an identical manner except that no partner was placed in the conditioning compartment. To remove odor, bedding was replaced after each session. The amount of time each mouse spent in the two compartments was measured during the 30-min pre- and postconditioning test sessions using EthoVision XT 10 (Noldus Information Technology, Wageningen, the Netherlands).

### Statistical analysis

We used statistical analytical software (GraphPad Prism 9.4.1 and SPSS 28) to analyze our data by analysis of variance (ANOVA) for more than two groups or Student’s t-test for two groups. The normality of data was evaluated by Shapiro–Wilk tests and the homogeneity of variance was evaluated by Levene’s test. When either assumption was violated, data were analyzed by Mann– Whitney test for unpaired data and Wilcoxon nonparametric test for paired data. The minimum level of significance was set a P<0.05. When multiple tests were applied in a data set, the level of significance was adjusted by Benjamini-Hochberg correction with a false-discovery rate of 5%, 10%, and 25%.

## RESULTS

### *Tbx1* deficiency alters the volumes of focal regions

We applied volumetric MRI to determine the volumes of the whole brain and brain regions of adult *Tbx1* HT mice and their wild-type (WT) littermates. We chose an *ex vivo* preparation, as it allows a longer scanning time and more accurate, reproducible data than *in vivo* scanning (50).

The volume of the whole brain was found to be unaffected by *Tbx1* heterozygosity (**Fig. S1**). We next applied a volumetric analysis to 19 atlas-based regions (**Fig. S2**) (45). A significant difference between WT and HT mice was found in the percentage values of the amygdala size relative to the whole brain (**Table S1; Fig. 1AB**), although this significance did not survive correction for multiple comparisons. The amygdala was the only region with an effect size larger than 1.0 (D=-1.18515), and the fimbria had the next-largest effect size (D=0.952486; **Table S1**). Eight regions had a nonsignificant effect size (|D|<0.2), and the remaining ten regions had small effect sizes (|D|>0.2) (see **Table S1**).

**Figure 1.**
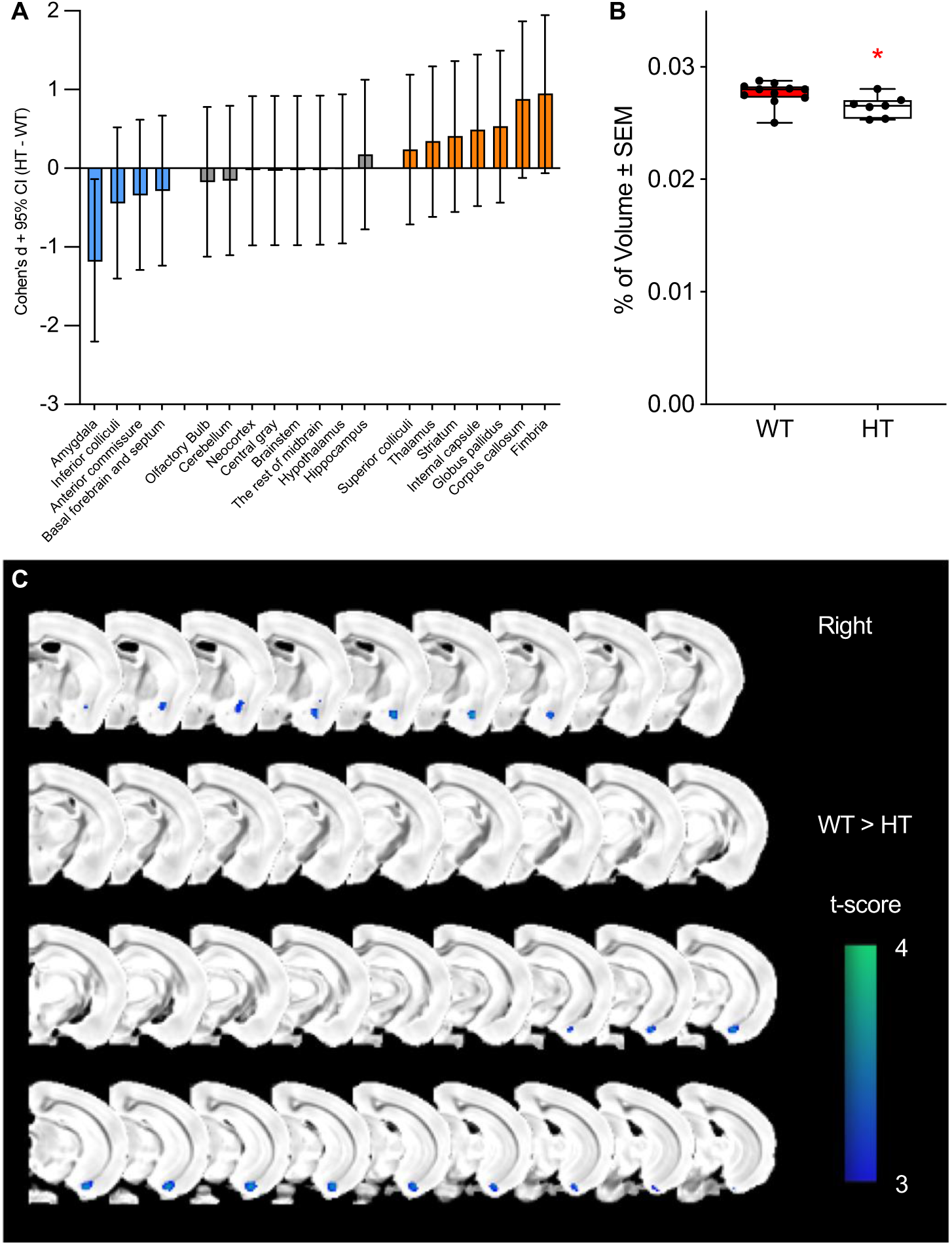
(**A**) Cohen’s D values of the % volume differences between wild-type (WT) and *Tbx1* heterozygous (HT) mice. The % volume of each region relative to the total volume was used to compute Cohen’s D values (HT−WT). In HT mice, volume increases are shown as positive values, and volume decreases are shown as negative values (see **Table S1** for all values). WT, n=11; HT, n=7. CI, confidence interval. (**B**) The mean % ± standard error of the mean of the amygdala relative to the whole brain volume (mm^3^) is smaller in heterozygous (HT) mice than in wild-type (WT) mice (t(16)=2.451, p=0.026). WT, n=11; HT, n=7. *p<0.05. (**C**) Voxel-based analysis of two-group differences between WT and HT mice. Permutation-based nonparametric testing with 500 random permutations was performed (small volume correction within the amygdala region of interest, p<0.05). A total of 409 voxels (0.21 mm^3^) survived this testing.

The amygdala, as defined by the mouse brain atlas of Ma and colleagues (45), is prolonged in the anteroposterior dimension and contains heterogeneous nuclei (**Fig. S3**). We thus applied a voxel analysis to detect focal volume alterations within the amygdala. The smaller amygdala volumes were detected in the anterior and posterior, but not middle, subregions of the amygdala (**Fig. 1C**).

While the neocortex as a whole (see **Fig. S2**) was not significantly altered in its volume (**Table S1**), it also is highly heterogeneous. In fact, individuals with 22q11.2 hemizygous deletions have thicker cortical gray matter and focal thickness reduction in restricted areas, such as the temporal and cingulate cortices (14). The volumes of various regions of the frontal cortex were reported to be either increased or reduced in a mouse model of 22q11.2 hemizygous deletion (16). Thus, we next analyzed the volumes of various cortical segments relative to the whole-brain volume. None of the 72 cortical and related regions were statistically significantly different in volume compared with WT mice after correction for multiple comparisons (**Table S2**). However, 10 brain regions had Cohen’s D values greater than 1.0 or smaller than −1.0 (see **Table S2**, red Cohen’s D values). When these 10 selected regions were analyzed as a set, their statistically significant differences survived correction for multiple comparisons (**Table S2**; **Fig. 2A**). Volume increases were found in the temporal association area, ventral area of the secondary auditory cortex, and primary auditory cortex of HT mice. Seven brain regions in HT mice had volume decreases: the ventral intermediate entorhinal cortex, amygdalopiriform transition area, secondary motor cortex, primary somatosensory cortex (jaw regions), medial entorhinal cortex, posteromedial cortical amygdaloid area, and cortex-amygdaloid transition zones.

**Figure 2.**
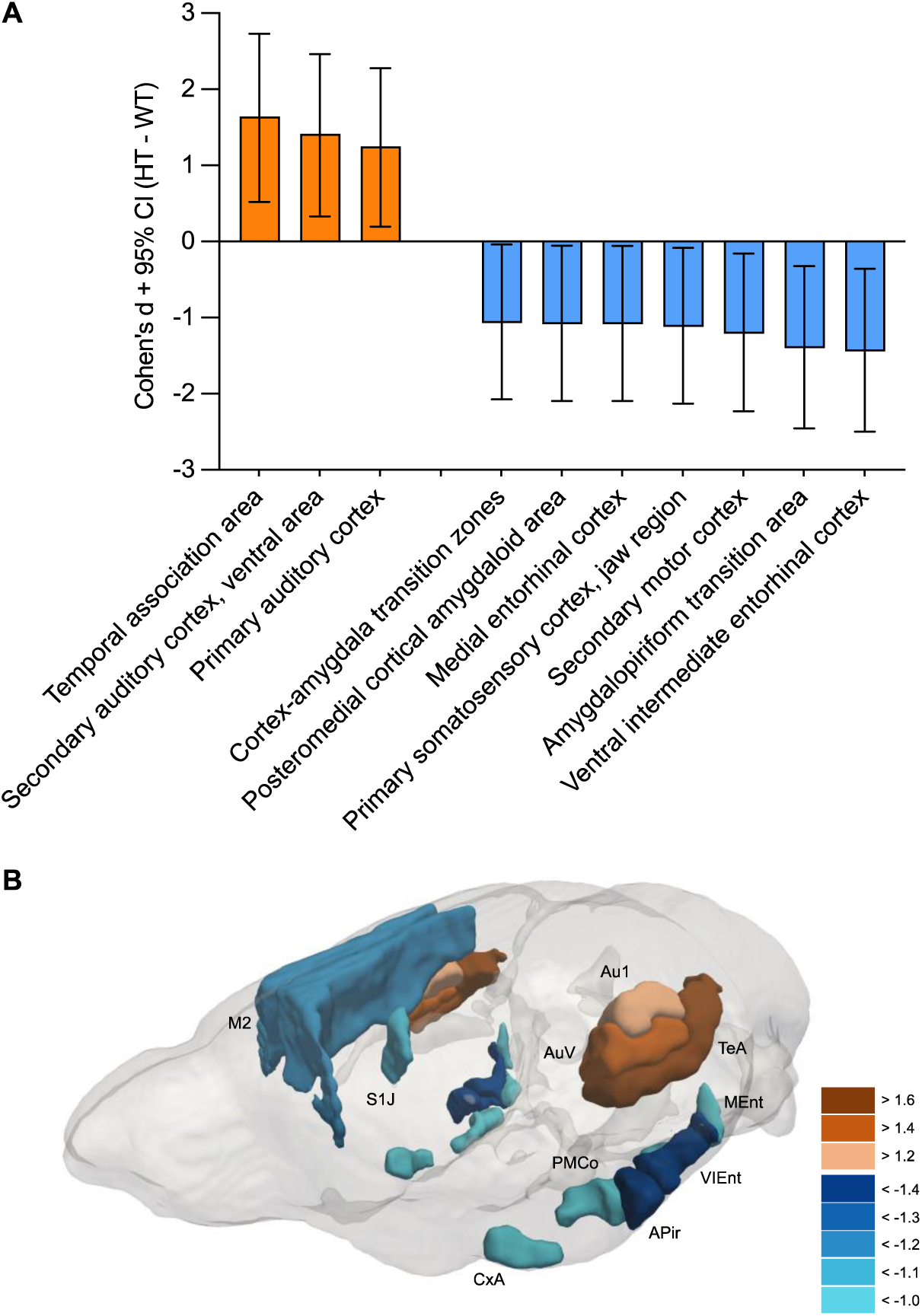
(**A**) Cohen’s D values of the percent volume differences between wild-type (WT) and *Tbx1* heterozygous (HT) mice. The percent volume of each cortical segment relative to the total volume was used to compute Cohen’s D values (HT−WT). In HT mice, volume increases are shown as positive values, and volume decreases are shown as negative values (see **Table S2** for all values). Among 72 cortical regions examined (see **Table S2**), only those with Cohen’s D values greater than 1.0 and smaller than −1.0 are listed. The statistically significant differences between the selected 10 regions between WT and HT mice survived the correction for multiple comparisons with a false-discovery rate of 5%. WT, n=11; HT, n=7. CI, confidence interval. (**B**) A 3D visualization of cortical segments with Cohen’s D values greater than 1.0 and smaller than −1.0. A clustering of volume reductions in and around the amygdala is noticeable (see also **Fig. S4** for a movie version).

A 3D image of regions with significant effect sizes indicated that cortical volume reductions clustered around the amygdala and that volume increases occurred in the auditory region and its surrounding cortical regions (**Fig. 2B; Fig. S4**). A cluster of regions with volume decreases in the posteroventral portion of the brain (**Fig. 2B; S4**) included the ventral intermediate entorhinal cortex, amygdalopiriform transition area, medial entorhinal cortex, posteromedial cortical amygdaloid area, and cortex amygdaloid transition zones, and these regions are all inside the boundary of the atlas-based amygdala (see **Fig. 2B; S4**).

Our ChIP-seq (chromatin immunoprecipitation followed by sequencing) data identified TBX1 binding sites near genes relevant to inhibitory neurotransmission, including *Gad2*, *Cep112*, *Gphn*, *Nlgn1*, *Nlgn2*, *Nlgn3*, *Nlgn4X*, and *Nlgn4Y* (51). As calretinin (also known as calbindin 2) mRNA colocalizes with a fraction of GABAergic neurons in the rodent amygdala (52) and is enriched in the amygdalopiriform transition zone (53), we evaluated by immunohistochemistry the size of the calretinin-positive neuropil in that region. The calretinin-rich neuropil of this region was significantly smaller in *Tbx1* HT mice than in their WT littermates (**Fig. 3A, B**).

**Figure 3.**
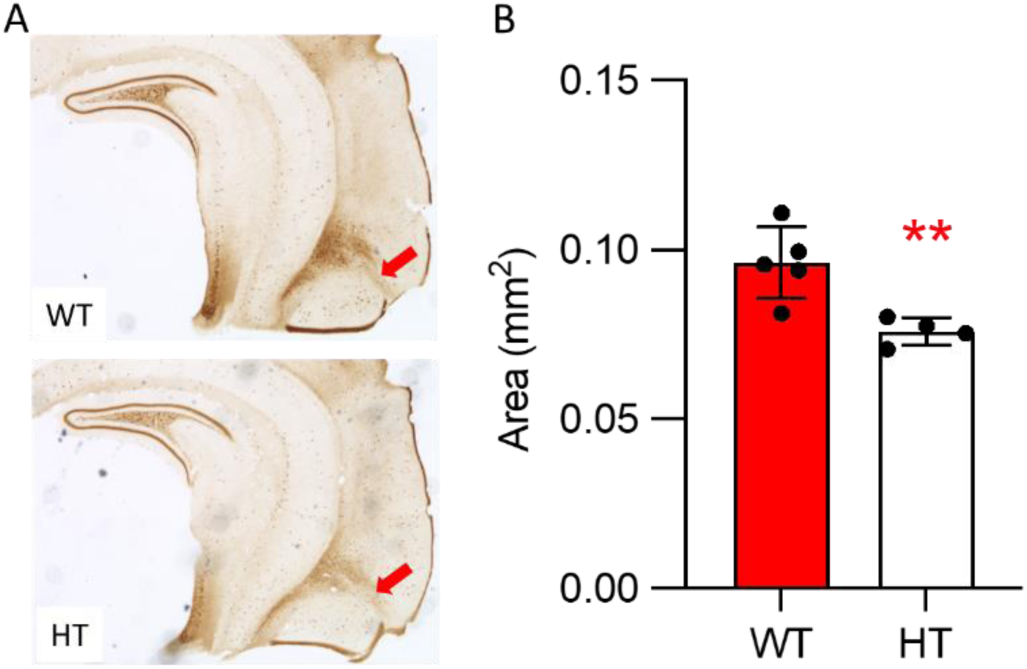
**(A)** Calretinin-positive neuropil in the amygdalopiriform transition area (red arrows) in wild-type (WT) and *Tbx1* heterozygous (HT) mice. The calretinin-positive neuropil was delineated within the anteroposterior extent of the region (Bregma from −2.46 mm to −3.64 mm from bregma), WT, n=5; HT, n=4. **(B)** The size of the calretinin-enriched neuropil in the amygdalopiriform transition zone was smaller in HT than in WT mice (t(7)=3.586, **p=0.0089).

### *Tbx1* deficiency impairs incentive values of a social partner

As the amygdala complex and its surrounding cortical regions showed robust volume reductions in *Tbx1* HT mice, we capitalized on a place-conditioning procedure, a classical conditioning that is known to rely on the amygdala. The incentive values of a littermate partner can be objectively evaluated by pairing a mouse with a partner in the presence of a specific set of proximal cues in a compartment, and a conditioned preference for that compartment can then be measured without a social partner. Despite the connotation of “place” conditioning, it has been established that conditioning with natural or drug-induced incentive stimuli is impaired in this procedure by damage to the amygdala, but is unaffected—or rather improved—by damage to the fimbria/fornix (54, 55, 56, 57, 58).

Each mouse was paired with either a littermate of the same genotype (*Tbx1* WT and *Tbx1* WT pair, or *Tbx1* HT and *Tbx1* HT pair) or of a different genotype (*Tbx1* WT and *Tbx1* HT pair, or *Tbx1* HT and *Tbx1* WT pair) in their home cages (**Fig. 4A**). After evaluating the baseline of their preference or aversion for the two compartments in the apparatus (i.e., preconditioning test, Day 1), a mouse was confined, with the center door closed, to one of the two compartments (i.e., paper bedding) with a social littermate partner for 24 hrs (conditioning, Day 2). On the next day, the subject mouse was confined, without a social littermate partner, in the other compartment (Cob bedding) for another 24 hrs (conditioning, Day 3). On the fourth day, each subject mouse was tested for their conditioned preference for the two compartments in the absence of the social partner (i.e., postconditioning test, Day 4).

**Figure 4.**
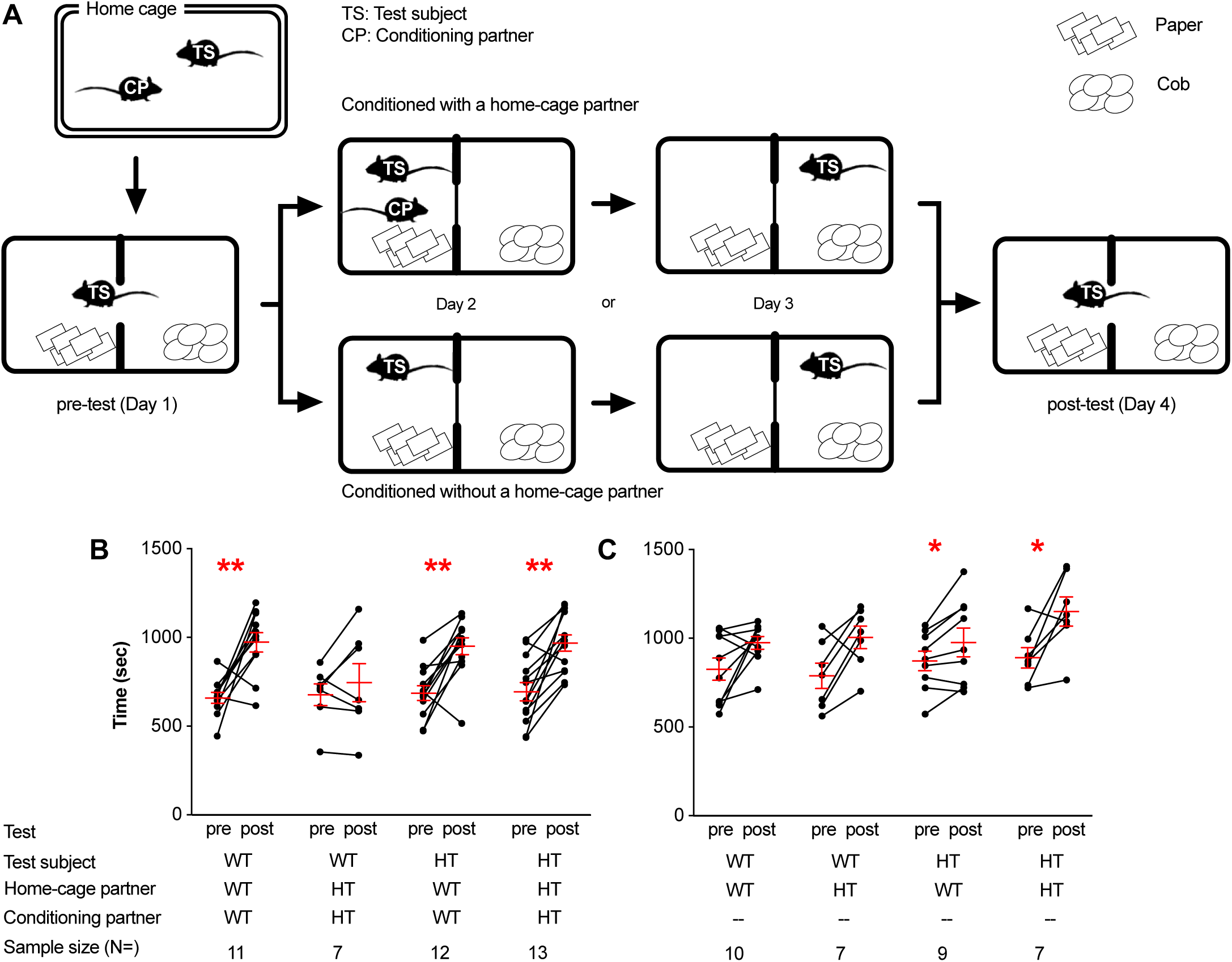
**A**) Experimental design. Each mouse was given a 30-min preconditioning session, two 24-hr conditioning sessions, and a 30-min postconditioning session. Each mouse test subject was caged with a littermate cage partner of the same or different genotype from weaning to testing. During conditioning, each test subject was conditioned with its own cage partner in one compartment and no partner in the other. Mice from a separate group were placed in both compartments without their cage partner. On the postconditioning day, each mouse was evaluated for their preference for or avoidance of each compartment without the cage partner. The average ± standard error of the mean time shift was calculated from the pre- to postconditioning sessions of wild-type (WT) and *Tbx1* heterozygous (HT) mice, conditioned with (**B**) or without (**C**) a home-cage partner. Wilcoxon nonparametric tests were applied to examine a shift in time from preconditioning session to postconditioning session. *p<0.05, **p<0.01.

*Tbx1* WT mice had a preference for a compartment where they were previously conditioned with a *Tbx1* WT littermate (**Fig. 4B**, WT/WT/WT). In a control experiment in which *Tbx1* WT mice were not conditioned in the apparatus, no significant shift appeared (**Fig. 4C**, WT/WT/--), indicating that the partner-dependent preference is not due to a spontaneous, partner-independent preference shift. In contrast, when conditioned with a *Tbx1* HT littermate, *Tbx1* WT mice did not exhibit a conditioned preference (**Fig. 4B**, WT/HT/HT); no shift in preference was also seen when no conditioning with a social partner was given (**Fig. 4C**, WT/HT/--), indicating that WT mice did not perceive the positive incentive value in HT partners.

*Tbx1* HT mice equally showed significant preferences after conditioning with either a *Tbx1* WT (**Fig. 4B**, HT/WT/WT) or *Tbx1* HT partner (**Fig. 4B**, HT/HT/HT). However, this was not due to a partner-dependent shift. When *Tbx1* HT mice were not conditioned with any mouse in the conditioning apparatus, they still showed an equally significant shift in preference (**Fig. 4C**, HT/WT/-- and HT/HT/--). In other words, the apparent conditioned preference for a compartment where HT mice were conditioned with a social partner was not significantly different from a spontaneous shift in preference without conditioning with a social partner.

*Tbx1* HT mice had a spontaneous shift to a compartment when the first and second conditioning sessions without a partner session were performed 48 and 24 hrs (Day 2 and Day 3), respectively, before testing on Day 4. Because of the longer temporal distance (i.e. less memory strength), the compartment of the first conditioning session (Day 2) might be perceived as more novel than the compartment of the second conditioning session (Day 3). This apparent preference for a more novel compartment in HT mice is consistent with our previous observation that *Tbx1* HT mice show a higher level of exploration of a novel object compared with WT mice (31).

## DISCUSSION

There are some inconsistencies among human imaging data of patients with 22q11.2 deletion (59). This is likely due to the unavoidably small number of samples of cases with rare genetic variants (9, 10). While analysis of 22q11.2 hemizygous carriers or its mouse models does not pinpoint the impacts of single genes, there are ultrarare cases of *TBX1* variants without 22q11.2 deletions with neurodevelopmental disorders (5, 21, 22, 23, 60). However, their neuronal or cognitive dimensions are not known, and their very rare nature does not permit imaging with a large sample size. Our data show that constitutive *Tbx1* heterozygosity results in highly focal volume alterations in anterior and posterior parts of the amygdaloid complex and its surrounding cortical regions, as is seen in 22q11.2 deletion carriers with psychosis (15) or ASD diagnosis (7) and in a genetic mouse model of 22q11.2 hemizygous deletion (16). These structural alterations coexisted with reduced sensitivity to a social partner in an amygdala-dependent task. Our mouse data predict changes in behavioral and structural dimensions in carriers of loss-of-function *TBX1* variants in humans and provide targets and readouts that could be evaluated for their therapeutic efficacy for carriers of variants of *TBX1* and 22q11.2 deletion.

Although a volume reduction in the atlas-based amygdaloid complex region of HT mice was statistically significant by itself, it failed to survive correction for multiple comparisons when all regions were analyzed as a set. However, three lines of evidence indicate that this is not a false-positive result. First, this impact of *Tbx1* deficiency did not follow the atlas-based boundary of the amygdaloid complex and was highly restricted to subregions of the amygdala. Our voxel-based analysis showed a volume reduction in HT mice in only the parts of the amygdala anterior to bregma −1.00 and posterior to bregma −2.70 (see **Fig. 1C**). Second, our analysis of cortical segments identified many regions with large effect sizes (D≤−1.0 and D≥1.0) inside and around the boundary of the atlas-based amygdaloid complex area: the cortex amygdaloid transition zone in the anterior subregion of the amygdaloid complex, and the amygdalopiriform transition area, ventral intermediate entorhinal cortex, posteromedial cortical amygdaloid area, and medial entorhinal cortex in the posterior subregion (see **Fig. 2**). A previous report showed a mouse model of 22q11.2 hemizygosity similarly did not display an overall volume alteration in the amygdala, but a voxel-wise measurement identified a statistically significant volume reduction in the anterior subregion of the amygdala (16). Finally, our immunohistochemical analysis validated a volume reduction in the amygdalopiriform transition area.

Our atlas-based analysis is likely to be compromised by two factors: an effect on specific subregions within the atlas-based amygdala area and simultaneous analysis of many regions with no alterations. The atlas-based analysis might lose the capacity to detect significant volume alterations within subregions of the entire amygdaloid complex. Moreover, in cases where specific brain regions are impacted, correction for multiple comparisons increases a false-negative rate. The initial failure to detect a significant volume reduction in the amygdala with the atlas-based MRI analysis is likely due to the highly focal impact of *Tbx1* deficiency on specific subregions of the anterior and posterior amygdala.

A corollary of our observation is that a conventional atlas-based anatomical classification might underestimate structural alterations. Moreover, statistical stringency might make it difficult to detect a focal, overall weak effect of one region among many examined regions. Supporting these possibilities, gene expression often does not follow the conventional atlas-based, histologically defined region classification; it can occur in only parts of a given region or can cut across the conventional border of two or more regions. Similarly, cell types based on gene-expression profiles often ignore the classical anatomical units (61, 62). Thus, a weak or nonsignificant effect following correction for multiple comparisons should be interpreted with caution to avoid false-negative cases.

We do not know why the impact of *Tbx1* deficiency is restricted to the anterior and posterior amygdaloid regions. One possibility is that there might be some specific time windows in which TBX1 deficiency impacts neurogenesis in distinct regions. Embryonic neurogenesis of these regions starts and peaks later than does cortical neurogenesis in neocortical regions in rodents (63, 64). Further supporting this possibility, *Tbx1* heterozygosity in stem cells initiated during the neonatal period is more effective in negatively impacting a later development of social interaction than that initiated in the same cell population postnatally (51). Moreover, *Tbx1* deficiency alters the embryonic neurogenesis of the primary somatosensory cortex but not that of the motor cortex (65).

In addition to the volume changes in these amygdaloid subregions, volumes were increased in the auditory cortical regions and decreased in the jaw region of the primary somatosensory cortex and the secondary motor cortex of *Tbx1* HT mice compared with their WT littermates. These data are consistent with a report that *Tbx1* deficiency results in thinning of the primary somatosensory cortex in adult mice due to alterations of embryonic neurogenesis (65); our data are, however, inconsistent with their observation that *Tbx1* deficiency has no impact on the primary and secondary motor cortices in adult mice with *Tbx1* deficiency (65). While the reason for this apparent discrepancy remains unclear, the study by Flore and colleagues used *Tbx1*^lacz/+^ mice with a C57BL/6N background (65), and our study used congenic *Tbx1*^+/−^ mice with a C57BL/6J background. These two C57BL/6 substrains carry non-identical alleles and result in phenotypic differences in gene expression, cellular, anatomical and behavioral phenotypes (17). *TBX1* deficiency results in craniofacial abnormalities, including cleft palate in mice (66, 67, 68, 69) and in humans (70) and hearing loss in mice (66) and sensorineural deafness in humans (21). These peripheral structural alterations might relate to volume changes in the corresponding regions in the brain (jaw region of the primary somatosensory cortex and the auditory cortex). It remains unclear if *Tbx1* deficiency simultaneously or sequentially impacts these brain regions and peripheral structures or vice versa. While more work is needed to untangle the causal links among the structures and functions that the gene variant impacts, our observations provide an opportunity to further understand the coordinated alterations in brain and peripheral structures.

Individuals with 22q11.2 hemizygous deletion have amygdala volume reductions (7, 15) and impaired activation of the right amygdala while processing the emotions of faces (71). A coisogenic mouse model of a large 22q11.2 hemizygous deletion is selectively impaired in an amygdala-dependent version of fear conditioning (i.e., another cue-dependent task) (72). *Tbx1* deficiency alone is sufficient to impair social incentive learning (current study), social interaction (31, 51), and neonatal social communication (30, 73) in mice. The amygdala is the site of action of another 22q11.2-encoded gene, *Sept5*, linked to social behavior (74). Taken together, amygdala-related behavioral phenotypes of 22q11.2 hemizygosity are, at least partly, likely to occur through *Tbx1* and *Sept5* deficiency. We do not exclude the possibility that other genes in 22q11.2 CNV might additionally or synergistically impact the amygdala. However, the currently available data emphasize the noncontiguous effects of 22q11.2 genes, in that not all genes are responsible for a given phenotype and that some gene products alone induce phenotypes (12, 13, 17, 18, 75, 76, 77).

Regions other than the amygdala complex are altered in size in carriers of 22q11.2 hemizygosity and in a mouse model of 22q11.2 hemizygosity. In humans with 22q11.2 hemizygous deletion, the caudate nucleus and nucleus accumbens are increased and the thalamus, putamen, hippocampus, and amygdala are reduced in volume (15). As the caudate nucleus, putamen, and nucleus accumbens are not separated and are collectively classified as the striatum in the rodent brain atlas, the opposing changes within this structure are likely to be difficult to detect in rodents.

Reported volume reductions in the hippocampus and thalamus in human carriers of 22q11.2 hemizygosity were not seen in our *Tbx1* HT mice (see **Fig. 1A**; **Table S1**). In a mouse model of 22q11.2 hemizygosity, the volume of the cerebellum is also reduced (16). We did not see a reduction in cerebellar volume in our mouse model of *Tbx1* heterozygosity. As far as TBX1-dependent causal chain from brain volume alterations to impaired social incentive learning is concerned, the cerebellum, hippocampus, thalamus, or striatum does not seem to be essential.

In conclusion, our mouse volumetric MRI study capitalized on the association of neurodevelopmental disorders and rare loss-of-function variants of *TBX1*, a 22q11.2 CNV-located gene, and provides a potential structural substrate for association between amygdala volume reductions and social incentive learning. A corollary of our observations is that therapeutic interventions for defective social incentive learning associated with constitutive *TBX1* deficiency need to be initiated at the developmental processes that determine the size of the amygdala and its surrounding cortical regions.

### Limitations

We selected specific brain regions and their functional outcomes based on an MRI screening. Imaging by MRI might lack the resolution to detect subtle alterations in the mouse brain. While a voxel analysis partially mitigated this interpretative limitation for the amygdala, it was not applied to all regions. Thus, our MRI data cannot indicate that the observed alterations are the only existing brain phenotypes of *Tbx1* deficiency.

We identified many other regions with medium to small effect sizes. While the importance of these regions could not be determined with sufficient statistical power, it is noteworthy that the fimbria sizes were increased and had the second-largest effect size (Cohen’s D=0.952486). This could be at least partly due to more myelination of medium-sized axons (although large myelinated axons are absent) in the fimbria of *Tbx1* HT mice (32). More work is needed to determine the causal roles of these structural alterations in various behavioral dimensions associated with *TBX1* deficiency and 22q11.2 CNV.

## Acknowledgments

The research reported in this publication was partly supported by the National Institutes of Health (R01MH099660, R01DC015776, R21HD053114, and R03HD108551) to N.H. The content is solely the responsibility of the authors and does not necessarily represent the official views of the National Institutes of Health. We thank Dr. Bernice Morrow for providing the original breeders of the *Tbx1* heterozygous mice.

## Author Contributions

T. Hiramoto and N. Hiroi designed the entire study and analyzed all data. A. Sumiyoshi, R. Ryoke, H. Nonaka, and R. Kawashima designed and performed the MRI study and analyzed data. R. Kato, A. Machida, K. Nomoto, K. Mogi, and T. Kikusui designed and performed the social conditioned preference experiment and analyzed data. G. Kang maintained and genotyped the mouse colony of *Tbx1* WT and HT mice. T. Yamauchi, B. Matsumura, Y.J. Hiroi performed immunohistochemistry of calretinin; B. Matsumura and Y.J. Hiroi analyzed data and prepared the related figures. T. Hiramoto, A. Sumiyoshi, R. Kato, T. Yamauchi, G. Kang, T. Kikusui, and N. Hiroi wrote the manuscript.

---

**Figure S1.**
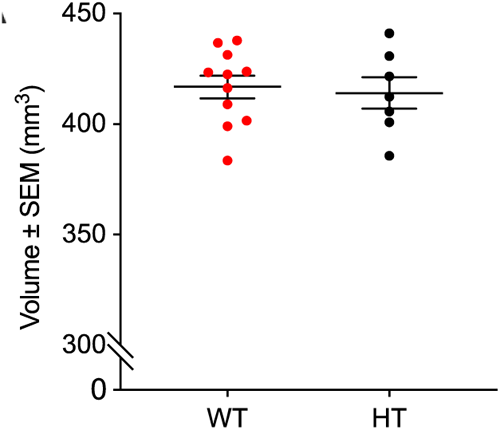
The mean ± standard error of the mean whole-brain volumes did not differ between wild-type (WT) and *Tbx1* heterozygous (HT) mice (t(16)=0.332, p=0.745). WT, n=11; HT, n=7.

**Figure S2.**
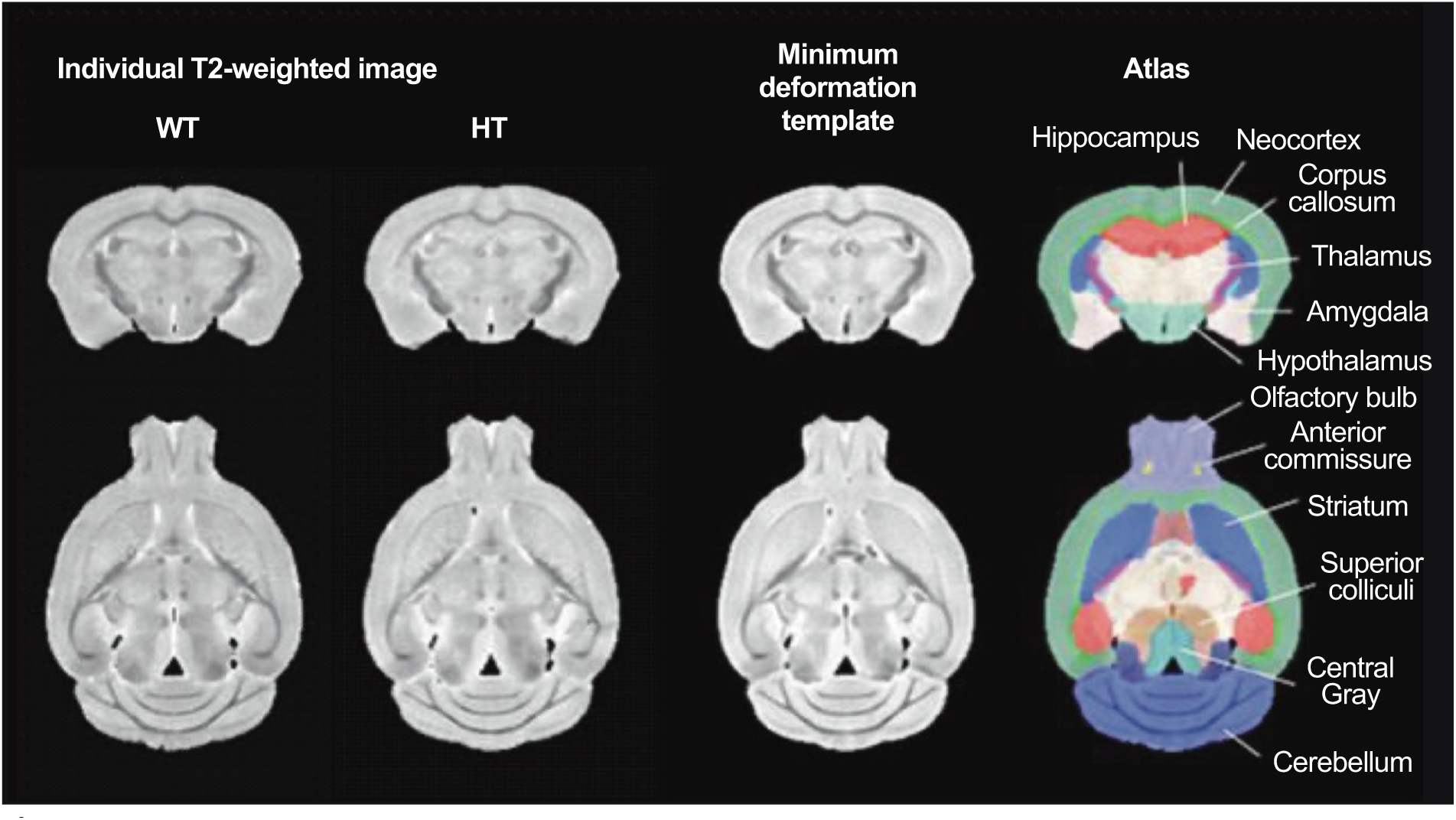
The brain regions used for analysis, following the regional classification of Ma and colleagues (45, 78). WT, wild-type; HT, heterozygous.

**Figure S3.**
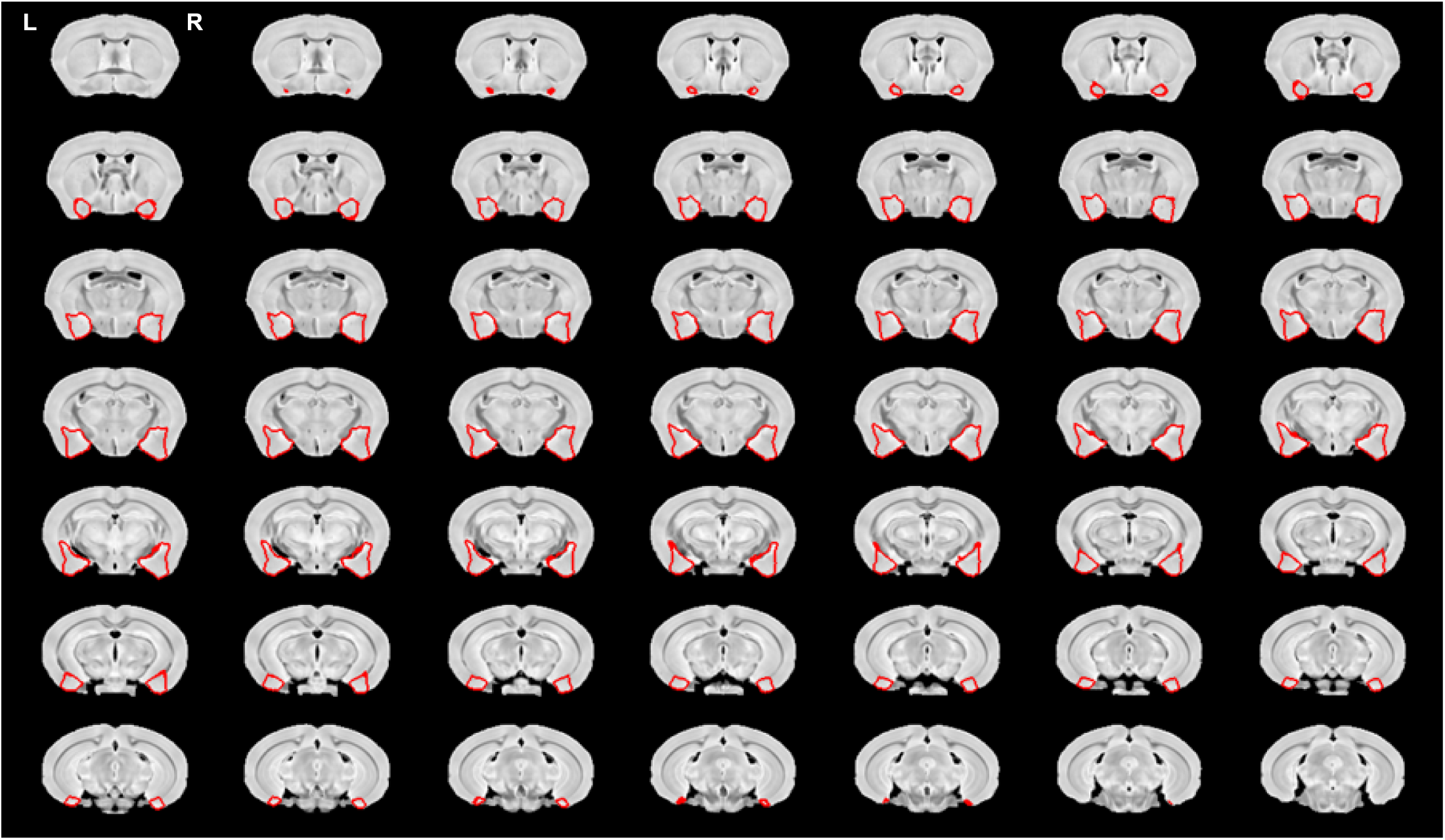
The location of the amygdala used for analysis. It includes the entire amygdaloid complex and its surrounding cortical regions as defined by a published template (45).

**Figure S4** A movie version of Figure 2B

**Table S1.**

| Regions | genotype | Average | s.e.m | T value | N | P value | Cohen's d |
| --- | --- | --- | --- | --- | --- | --- | --- |
| **HT < WT** |  |  |  |  |  |  |  |
| Amygdala | WT | 2.76E-02 | 3.08E-04 | -2.45E+00 | 16 | 2.61E-02 | -1.19E+00 |
|  | HT | 2.65E-02 | 3.59E-04 |  |  |  |  |
| Inferior colliculi | WT | 1.48E-02 | 1.99E-04 | -9.31E-01 | 16 | 3.66E-01 | -4.50E-01 |
|  | HT | 1.45E-02 | 2.32E-04 |  |  |  |  |
| Anterior commissure | WT | 1.57E-03 | 1.68E-05 | -7.15E-01 | 16 | 4.85E-01 | -3.46E-01 |
|  | HT | 1.55E-03 | 2.18E-05 |  |  |  |  |
| Basal forebrain and septum | WT | 3.01E-02 | 3.37E-04 | -6.01E-01 | 16 | 5.56E-01 | -2.91E-01 |
|  | HT | 2.97E-02 | 5.10E-04 |  |  |  |  |
| **HT = WT, D ≤ -0.2** |  |  |  |  |  |  |  |
| Olfactory Bulb | WT | 6.24E-02 | 1.04E-03 | -3.65E-01 | 16 | 7.20E-01 | -1.77E-01 |
|  | HT | 6.17E-02 | 1.34E-03 |  |  |  |  |
| Cerebellum | WT | 1.23E-01 | 1.03E-03 | -3.29E-01 | 16 | 7.46E-01 | -1.59E-01 |
|  | HT | 1.22E-01 | 1.91E-03 |  |  |  |  |
| Neocortex | WT | 3.14E-01 | 2.01E-03 | -5.93E-02 | 16 | 9.53E-01 | -2.87E-02 |
|  | HT | 3.13E-01 | 3.37E-03 |  |  |  |  |
| Central gray | WT | 1.17E-02 | 1.42E-04 | -7.01E-02 | 16 | 9.45E-01 | -3.39E-02 |
|  | HT | 1.17E-02 | 2.64E-04 |  |  |  |  |
| Brainstem | WT | 1.14E-01 | 1.04E-03 | -5.97E-02 | 16 | 9.53E-01 | -2.89E-02 |
|  | HT | 1.14E-01 | 1.17E-03 |  |  |  |  |
| Rest of midbrain | WT | 2.46E-02 | 1.91E-04 | -5.42E-02 | 16 | 9.57E-01 | -2.62E-02 |
|  | HT | 2.46E-02 | 5.47E-04 |  |  |  |  |
| Hypothalamus | WT | 2.65E-02 | 2.87E-04 | -1.49E-02 | 16 | 9.88E-01 | -7.19E-03 |
|  | HT | 2.65E-02 | 2.29E-04 |  |  |  |  |
| Hippocampus | WT | 6.10E-02 | 4.51E-04 | 3.66E-01 | 16 | 7.20E-01 | 1.77E-01 |
|  | HT | 6.13E-02 | 6.49E-04 |  |  |  |  |
| **HT > WT, D ≥ 0.2** |  |  |  |  |  |  |  |
| Superior colliculi | WT | 2.60E-02 | 3.07E-04 | 4.97E-01 | 16 | 6.26E-01 | 2.40E-01 |
|  | HT | 2.63E-02 | 5.31E-04 |  |  |  |  |
| Thalamus | WT | 4.94E-02 | 6.59E-04 | 7.10E-01 | 16 | 4.88E-01 | 3.43E-01 |
|  | HT | 5.03E-02 | 1.37E-03 |  |  |  |  |
| Striatum | WT | 6.34E-02 | 5.42E-04 | 8.46E-01 | 16 | 4.10E-01 | 4.09E-01 |
|  | HT | 6.41E-02 | 6.12E-04 |  |  |  |  |
| Internal capsule | WT | 5.19E-03 | 6.20E-05 | 1.01E+00 | 16 | 3.27E-01 | 4.89E-01 |
|  | HT | 5.30E-03 | 1.08E-04 |  |  |  |  |
| Globus pallidus | WT | 6.79E-03 | 6.34E-05 | 1.11E+00 | 16 | 2.85E-01 | 5.35E-01 |
|  | HT | 6.92E-03 | 1.14E-04 |  |  |  |  |
| Corpus callosum | WT | 3.05E-02 | 2.09E-04 | 1.82E+00 | 16 | 8.68E-02 | 8.82E-01 |
|  | HT | 3.11E-02 | 1.73E-04 |  |  |  |  |
| Fimbria | WT | 5.66E-03 | 8.66E-05 | 1.97E+00 | 16 | 6.64E-02 | 9.52E-01 |
|  | HT | 5.97E-03 | 1.40E-04 |  |  |  |  |

**Table S2.**
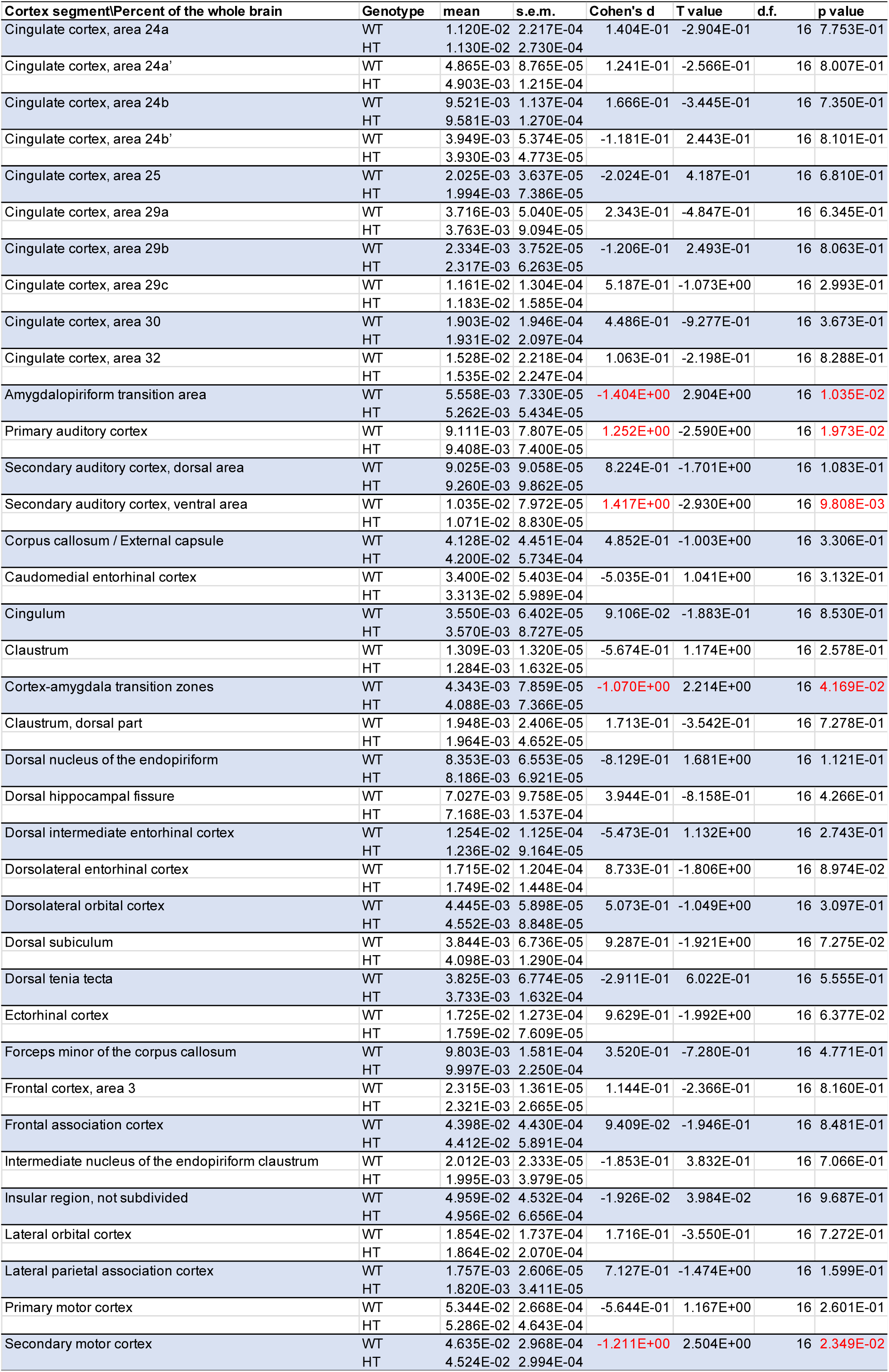
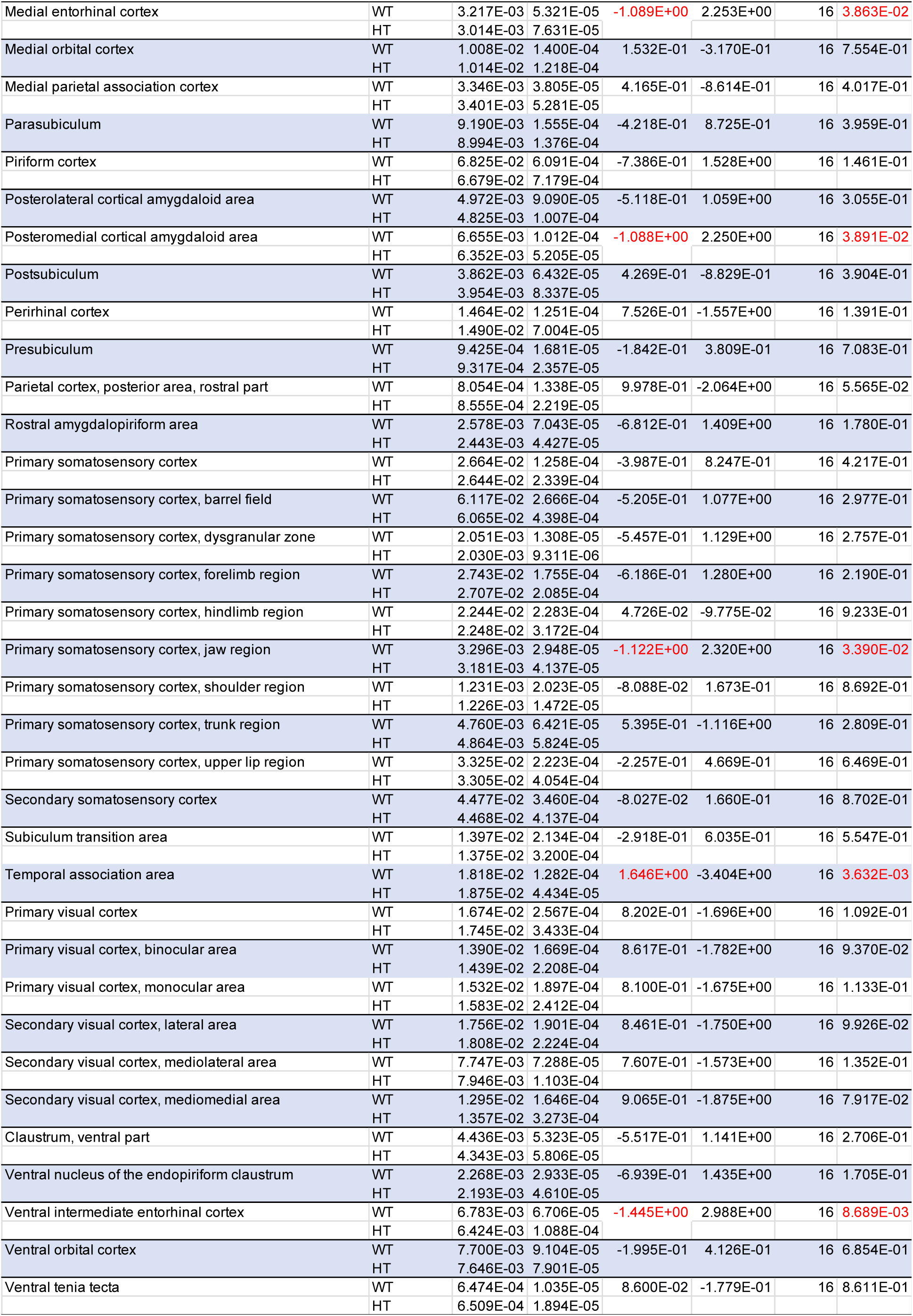

| Cortex segment | Percent of the whole brain | Genotype | mean | s.e.m. | Cohen's d | T value | d.f. | p value |
| --- | --- | --- | --- | --- | --- | --- | --- | --- |
| Cingulate cortex, area 24a |  | WT | 1.120E-02 | 2.217E-04 | 1.404E-01 | -2.904E-01 | 16 | 7.753E-01 |
|  |  | HT | 1.130E-02 | 2.730E-04 |  |  |  |  |
| Cingulate cortex, area 24a' |  | WT | 4.865E-03 | 8.765E-05 | 1.241E-01 | -2.566E-01 | 16 | 8.007E-01 |
|  |  | HT | 4.903E-03 | 1.215E-04 |  |  |  |  |
| Cingulate cortex, area 24b |  | WT | 9.521E-03 | 1.137E-04 | 1.666E-01 | -3.445E-01 | 16 | 7.350E-01 |
|  |  | HT | 9.581E-03 | 1.270E-04 |  |  |  |  |
| Cingulate cortex, area 24b' |  | WT | 3.949E-03 | 5.374E-05 | -1.181E-01 | 2.443E-01 | 16 | 8.101E-01 |
|  |  | HT | 3.930E-03 | 4.773E-05 |  |  |  |  |
| Cingulate cortex, area 25 |  | WT | 2.025E-03 | 3.637E-05 | -2.024E-01 | 4.187E-01 | 16 | 6.810E-01 |
|  |  | HT | 1.994E-03 | 7.386E-05 |  |  |  |  |
| Cingulate cortex, area 29a |  | WT | 3.716E-03 | 5.040E-05 | 2.343E-01 | -4.847E-01 | 16 | 6.345E-01 |
|  |  | HT | 3.763E-03 | 9.094E-05 |  |  |  |  |
| Cingulate cortex, area 29b |  | WT | 2.334E-03 | 3.752E-05 | -1.206E-01 | 2.493E-01 | 16 | 8.063E-01 |
|  |  | HT | 2.317E-03 | 6.263E-05 |  |  |  |  |
| Cingulate cortex, area 29c |  | WT | 1.161E-02 | 1.304E-04 | 5.187E-01 | -1.073E+00 | 16 | 2.993E-01 |
|  |  | HT | 1.183E-02 | 1.585E-04 |  |  |  |  |
| Cingulate cortex, area 30 |  | WT | 1.903E-02 | 1.946E-04 | 4.486E-01 | -9.277E-01 | 16 | 3.673E-01 |
|  |  | HT | 1.931E-02 | 2.097E-04 |  |  |  |  |
| Cingulate cortex, area 32 |  | WT | 1.528E-02 | 2.218E-04 | 1.063E-01 | -2.198E-01 | 16 | 8.288E-01 |
|  |  | HT | 1.535E-02 | 2.247E-04 |  |  |  |  |
| Amygdalopiriform transition area |  | WT | 5.558E-03 | 7.330E-05 | -1.404E+00 | 2.904E+00 | 16 | 1.035E-02 |
|  |  | HT | 5.262E-03 | 5.434E-05 |  |  |  |  |
| Primary auditory cortex |  | WT | 9.111E-03 | 7.807E-05 | 1.252E+00 | -2.590E+00 | 16 | 1.973E-02 |
|  |  | HT | 9.408E-03 | 7.400E-05 |  |  |  |  |
| Secondary auditory cortex, dorsal area |  | WT | 9.025E-03 | 9.058E-05 | 8.224E-01 | -1.701E+00 | 16 | 1.083E-01 |
|  |  | HT | 9.260E-03 | 9.862E-05 |  |  |  |  |
| Secondary auditory cortex, ventral area |  | WT | 1.035E-02 | 7.972E-05 | 1.417E+00 | -2.930E+00 | 16 | 9.808E-03 |
|  |  | HT | 1.071E-02 | 8.830E-05 |  |  |  |  |
| Corpus callosum / External capsule |  | WT | 4.128E-02 | 4.451E-04 | 4.852E-01 | -1.003E+00 | 16 | 3.306E-01 |
|  |  | HT | 4.200E-02 | 5.734E-04 |  |  |  |  |
| Caudomedial entorhinal cortex |  | WT | 3.400E-02 | 5.403E-04 | -5.035E-01 | 1.041E+00 | 16 | 3.132E-01 |
|  |  | HT | 3.313E-02 | 5.989E-04 |  |  |  |  |
| Cingulum |  | WT | 3.550E-03 | 6.402E-05 | 9.106E-02 | -1.883E-01 | 16 | 8.530E-01 |
|  |  | HT | 3.570E-03 | 8.727E-05 |  |  |  |  |
| Clastrum |  | WT | 1.309E-03 | 1.320E-05 | -5.674E-01 | 1.174E+00 | 16 | 2.578E-01 |
|  |  | HT | 1.284E-03 | 1.632E-05 |  |  |  |  |
| Cortex-amygdala transition zones |  | WT | 4.343E-03 | 7.859E-05 | -1.070E+00 | 2.214E+00 | 16 | 4.169E-02 |
|  |  | HT | 4.088E-03 | 7.366E-05 |  |  |  |  |
| Clastrum, dorsal part |  | WT | 1.948E-03 | 2.406E-05 | 1.713E-01 | -3.542E-01 | 16 | 7.278E-01 |
|  |  | HT | 1.964E-03 | 4.652E-05 |  |  |  |  |
| Dorsal nucleus of the endopiriform |  | WT | 8.353E-03 | 6.553E-05 | -8.129E-01 | 1.681E+00 | 16 | 1.121E-01 |
|  |  | HT | 8.186E-03 | 6.921E-05 |  |  |  |  |
| Dorsal hippocampal fissure |  | WT | 7.027E-03 | 9.758E-05 | 3.944E-01 | -8.158E-01 | 16 | 4.266E-01 |
|  |  | HT | 7.168E-03 | 1.537E-04 |  |  |  |  |
| Dorsal intermediate entorhinal cortex |  | WT | 1.254E-02 | 1.125E-04 | -5.473E-01 | 1.132E+00 | 16 | 2.743E-01 |
|  |  | HT | 1.236E-02 | 9.164E-05 |  |  |  |  |
| Dorsolateral entorhinal cortex |  | WT | 1.715E-02 | 1.204E-04 | 8.733E-01 | -1.806E+00 | 16 | 8.974E-02 |
|  |  | HT | 1.749E-02 | 1.448E-04 |  |  |  |  |
| Dorsolateral orbital cortex |  | WT | 4.445E-03 | 5.898E-05 | 5.073E-01 | -1.049E+00 | 16 | 3.097E-01 |
|  |  | HT | 4.552E-03 | 8.848E-05 |  |  |  |  |
| Dorsal subiculum |  | WT | 3.844E-03 | 6.736E-05 | 9.287E-01 | -1.921E+00 | 16 | 7.275E-02 |
|  |  | HT | 4.098E-03 | 1.290E-04 |  |  |  |  |
| Dorsal tenia tecta |  | WT | 3.825E-03 | 6.774E-05 | -2.911E-01 | 6.022E-01 | 16 | 5.555E-01 |
|  |  | HT | 3.733E-03 | 1.632E-04 |  |  |  |  |
| Ectorhinal cortex |  | WT | 1.725E-02 | 1.273E-04 | 9.629E-01 | -1.992E+00 | 16 | 6.377E-02 |
|  |  | HT | 1.759E-02 | 7.609E-05 |  |  |  |  |
| Forceps minor of the corpus callosum |  | WT | 9.803E-03 | 1.581E-04 | 3.520E-01 | -7.280E-01 | 16 | 4.771E-01 |
|  |  | HT | 9.997E-03 | 2.250E-04 |  |  |  |  |
| Frontal cortex, area 3 |  | WT | 2.315E-03 | 1.361E-05 | 1.144E-01 | -2.366E-01 | 16 | 8.160E-01 |
|  |  | HT | 2.321E-03 | 2.665E-05 |  |  |  |  |
| Frontal association cortex |  | WT | 4.398E-02 | 4.430E-04 | 9.409E-02 | -1.946E-01 | 16 | 8.481E-01 |
|  |  | HT | 4.412E-02 | 5.891E-04 |  |  |  |  |
| Intermediate nucleus of the endopiriform claustrum |  | WT | 2.012E-03 | 2.333E-05 | -1.853E-01 | 3.832E-01 | 16 | 7.066E-01 |
|  |  | HT | 1.995E-03 | 3.979E-05 |  |  |  |  |
| Insular region, not subdivided |  | WT | 4.959E-02 | 4.532E-04 | -1.926E-02 | 3.984E-02 | 16 | 9.687E-01 |
|  |  | HT | 4.956E-02 | 6.656E-04 |  |  |  |  |
| Lateral orbital cortex |  | WT | 1.854E-02 | 1.737E-04 | 1.716E-01 | -3.550E-01 | 16 | 7.272E-01 |
|  |  | HT | 1.864E-02 | 2.070E-04 |  |  |  |  |
| Lateral parietal association cortex |  | WT | 1.757E-03 | 2.606E-05 | 7.127E-01 | -1.474E+00 | 16 | 1.599E-01 |
|  |  | HT | 1.820E-03 | 3.411E-05 |  |  |  |  |
| Primary motor cortex |  | WT | 5.344E-02 | 2.668E-04 | -5.644E-01 | 1.167E+00 | 16 | 2.601E-01 |
|  |  | HT | 5.286E-02 | 4.643E-04 |  |  |  |  |
| Secondary motor cortex |  | WT | 4.635E-02 | 2.968E-04 | -1.211E+00 | 2.504E+00 | 16 | 2.349E-02 |
|  |  | HT | 4.524E-02 | 2.994E-04 |  |  |  |  |

Table S2 Continued
|  |  |  |  |  |  |  |  |
| --- | --- | --- | --- | --- | --- | --- | --- |
| Medial entorhinal cortex | WT | 3.217E-03 | 5.321E-05 | -1.089E+00 | 2.253E+00 | 16 | 3.863E-02 |
|  | HT | 3.014E-03 | 7.631E-05 |  |  |  |  |
| Medial orbital cortex | WT | 1.008E-02 | 1.400E-04 | 1.532E-01 | -3.170E-01 | 16 | 7.554E-01 |
|  | HT | 1.014E-02 | 1.218E-04 |  |  |  |  |
| Medial parietal association cortex | WT | 3.346E-03 | 3.805E-05 | 4.165E-01 | -8.614E-01 | 16 | 4.017E-01 |
|  | HT | 3.401E-03 | 5.281E-05 |  |  |  |  |
| Parasubiculum | WT | 9.190E-03 | 1.555E-04 | -4.218E-01 | 8.725E-01 | 16 | 3.959E-01 |
|  | HT | 8.994E-03 | 1.376E-04 |  |  |  |  |
| Piriform cortex | WT | 6.825E-02 | 6.091E-04 | -7.386E-01 | 1.528E+00 | 16 | 1.461E-01 |
|  | HT | 6.679E-02 | 7.179E-04 |  |  |  |  |
| Posterolateral cortical amygdaloid area | WT | 4.972E-03 | 9.090E-05 | -5.118E-01 | 1.059E+00 | 16 | 3.055E-01 |
|  | HT | 4.825E-03 | 1.007E-04 |  |  |  |  |
| Posteromedial cortical amygdaloid area | WT | 6.655E-03 | 1.012E-04 | -1.088E+00 | 2.250E+00 | 16 | 3.891E-02 |
|  | HT | 6.352E-03 | 5.205E-05 |  |  |  |  |
| Postsubiculum | WT | 3.862E-03 | 6.432E-05 | 4.269E-01 | -8.829E-01 | 16 | 3.904E-01 |
|  | HT | 3.954E-03 | 8.337E-05 |  |  |  |  |
| Perirhinal cortex | WT | 1.464E-02 | 1.251E-04 | 7.526E-01 | -1.557E+00 | 16 | 1.391E-01 |
|  | HT | 1.490E-02 | 7.004E-05 |  |  |  |  |
| Presubiculum | WT | 9.425E-04 | 1.681E-05 | -1.842E-01 | 3.809E-01 | 16 | 7.083E-01 |
|  | HT | 9.317E-04 | 2.357E-05 |  |  |  |  |
| Parietal cortex, posterior area, rostral part | WT | 8.054E-04 | 1.338E-05 | 9.978E-01 | -2.064E+00 | 16 | 5.565E-02 |
|  | HT | 8.555E-04 | 2.219E-05 |  |  |  |  |
| Rostral amygdalopiriform area | WT | 2.578E-03 | 7.043E-05 | -6.812E-01 | 1.409E+00 | 16 | 1.780E-01 |
|  | HT | 2.443E-03 | 4.427E-05 |  |  |  |  |
| Primary somatosensory cortex | WT | 2.664E-02 | 1.258E-04 | -3.987E-01 | 8.247E-01 | 16 | 4.217E-01 |
|  | HT | 2.644E-02 | 2.339E-04 |  |  |  |  |
| Primary somatosensory cortex, barrel field | WT | 6.117E-02 | 2.666E-04 | -5.205E-01 | 1.077E+00 | 16 | 2.977E-01 |
|  | HT | 6.065E-02 | 4.398E-04 |  |  |  |  |
| Primary somatosensory cortex, dysgranular zone | WT | 2.051E-03 | 1.308E-05 | -5.457E-01 | 1.129E+00 | 16 | 2.757E-01 |
|  | HT | 2.030E-03 | 9.311E-06 |  |  |  |  |
| Primary somatosensory cortex, forelimb region | WT | 2.743E-02 | 1.755E-04 | -6.186E-01 | 1.280E+00 | 16 | 2.190E-01 |
|  | HT | 2.707E-02 | 2.085E-04 |  |  |  |  |
| Primary somatosensory cortex, hindlimb region | WT | 2.244E-02 | 2.283E-04 | 4.726E-02 | -9.775E-02 | 16 | 9.233E-01 |
|  | HT | 2.248E-02 | 3.172E-04 |  |  |  |  |
| Primary somatosensory cortex, jaw region | WT | 3.296E-03 | 2.948E-05 | -1.122E+00 | 2.320E+00 | 16 | 3.390E-02 |
|  | HT | 3.181E-03 | 4.137E-05 |  |  |  |  |
| Primary somatosensory cortex, shoulder region | WT | 1.231E-03 | 2.023E-05 | -8.088E-02 | 1.673E-01 | 16 | 8.692E-01 |
|  | HT | 1.226E-03 | 1.472E-05 |  |  |  |  |
| Primary somatosensory cortex, trunk region | WT | 4.760E-03 | 6.421E-05 | 5.395E-01 | -1.116E+00 | 16 | 2.809E-01 |
|  | HT | 4.864E-03 | 5.824E-05 |  |  |  |  |
| Primary somatosensory cortex, upper lip region | WT | 3.325E-02 | 2.223E-04 | -2.257E-01 | 4.669E-01 | 16 | 6.469E-01 |
|  | HT | 3.305E-02 | 4.054E-04 |  |  |  |  |
| Secondary somatosensory cortex | WT | 4.477E-02 | 3.460E-04 | -8.027E-02 | 1.660E-01 | 16 | 8.702E-01 |
|  | HT | 4.468E-02 | 4.137E-04 |  |  |  |  |
| Subiculum transition area | WT | 1.397E-02 | 2.134E-04 | -2.918E-01 | 6.035E-01 | 16 | 5.547E-01 |
|  | HT | 1.375E-02 | 3.200E-04 |  |  |  |  |
| Temporal association area | WT | 1.818E-02 | 1.282E-04 | 1.646E+00 | -3.404E+00 | 16 | 3.632E-03 |
|  | HT | 1.875E-02 | 4.434E-05 |  |  |  |  |
| Primary visual cortex | WT | 1.674E-02 | 2.567E-04 | 8.202E-01 | -1.696E+00 | 16 | 1.092E-01 |
|  | HT | 1.745E-02 | 3.433E-04 |  |  |  |  |
| Primary visual cortex, binocular area | WT | 1.390E-02 | 1.669E-04 | 8.617E-01 | -1.782E+00 | 16 | 9.370E-02 |
|  | HT | 1.439E-02 | 2.208E-04 |  |  |  |  |
| Primary visual cortex, monocular area | WT | 1.532E-02 | 1.897E-04 | 8.100E-01 | -1.675E+00 | 16 | 1.133E-01 |
|  | HT | 1.583E-02 | 2.412E-04 |  |  |  |  |
| Secondary visual cortex, lateral area | WT | 1.756E-02 | 1.901E-04 | 8.461E-01 | -1.750E+00 | 16 | 9.926E-02 |
|  | HT | 1.808E-02 | 2.224E-04 |  |  |  |  |
| Secondary visual cortex, mediolateral area | WT | 7.747E-03 | 7.288E-05 | 7.607E-01 | -1.573E+00 | 16 | 1.352E-01 |
|  | HT | 7.946E-03 | 1.103E-04 |  |  |  |  |
| Secondary visual cortex, mediomedial area | WT | 1.295E-02 | 1.646E-04 | 9.065E-01 | -1.875E+00 | 16 | 7.917E-02 |
|  | HT | 1.357E-02 | 3.273E-04 |  |  |  |  |
| Clastrum, ventral part | WT | 4.436E-03 | 5.323E-05 | -5.517E-01 | 1.141E+00 | 16 | 2.706E-01 |
|  | HT | 4.343E-03 | 5.806E-05 |  |  |  |  |
| Ventral nucleus of the endopiriform claustrum | WT | 2.268E-03 | 2.933E-05 | -6.939E-01 | 1.435E+00 | 16 | 1.705E-01 |
|  | HT | 2.193E-03 | 4.610E-05 |  |  |  |  |
| Ventral intermediate entorhinal cortex | WT | 6.783E-03 | 6.706E-05 | -1.445E+00 | 2.988E+00 | 16 | 8.689E-03 |
|  | HT | 6.424E-03 | 1.088E-04 |  |  |  |  |
| Ventral orbital cortex | WT | 7.700E-03 | 9.104E-05 | -1.995E-01 | 4.126E-01 | 16 | 6.854E-01 |
|  | HT | 7.646E-03 | 7.901E-05 |  |  |  |  |
| Ventral tenia tecta | WT | 6.474E-04 | 1.035E-05 | 8.600E-02 | -1.779E-01 | 16 | 8.611E-01 |
|  | HT | 6.509E-04 | 1.894E-05 |  |  |  |  |

