## Supplementary figures and images for "Structural alterations in the amygdala and impaired social incentive learning in a mouse model of a genetic variant associated with neurodevelopmental disorders"

### Figure S4

#### Slide 1
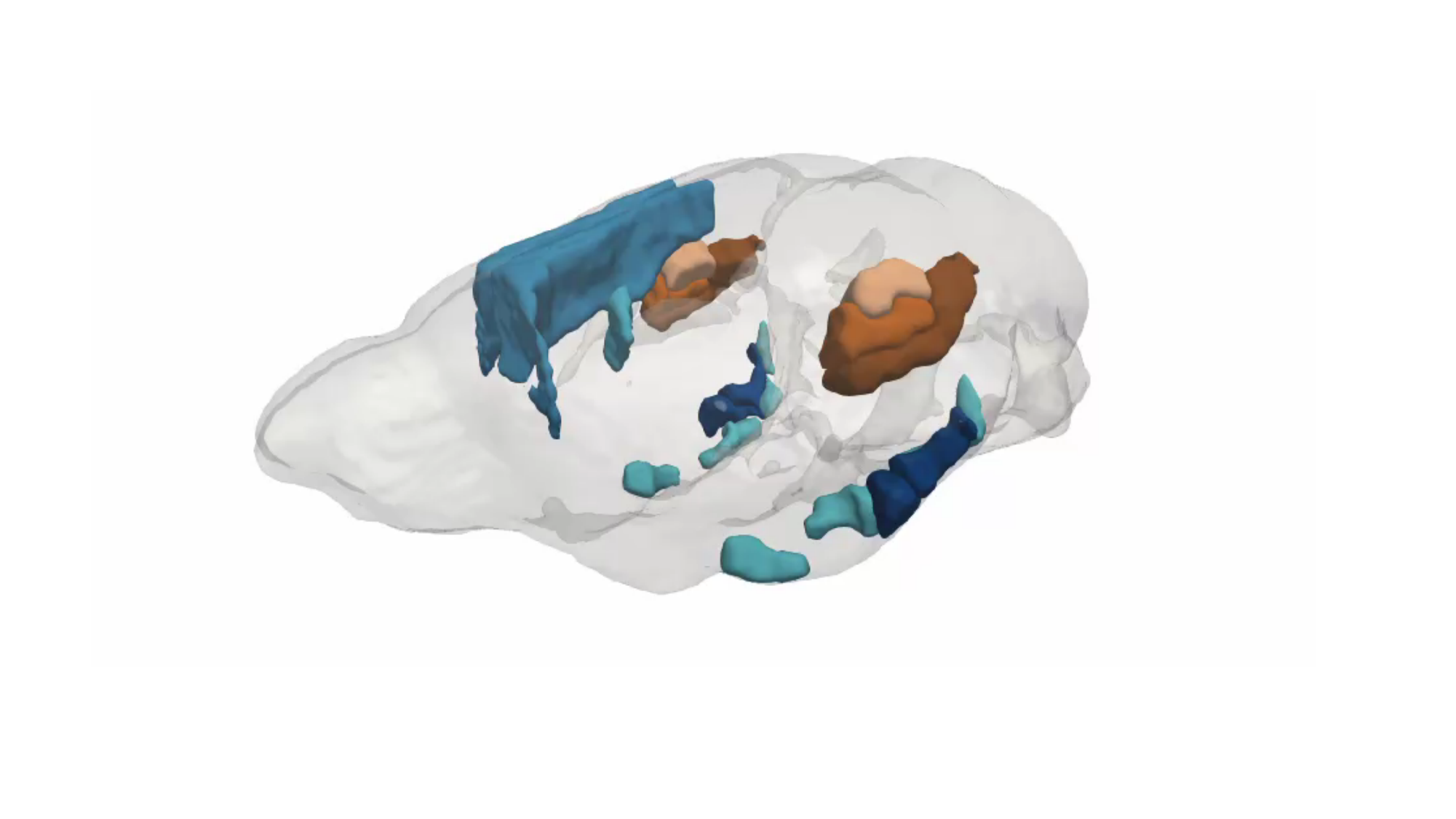
